# A *Clostridia*-rich microbiota contributes to increased excretion of bile acids in diarrhea-predominant irritable bowel syndrome

**DOI:** 10.1101/630350

**Authors:** Ling Zhao, Wei Yang, Yang Chen, Fengjie Huang, Lin Lu, Chengyuan Lin, Tao Huang, Ziwan Ning, Lixiang Zhai, Linda LD Zhong, Waiching Lam, Zhen Yang, Xuan Zhang, Chungwah Cheng, Lijuan Han, Qinwei Qiu, Xiaoxiao Shang, Runyue Huang, Zhenxing Ren, Dongfeng Chen, Silong Sun, Hani El-Nezami, Aiping Lu, Xiaodong Fang, Wei Jia, Zhaoxiang Bian

**Author notes:** Authors share co-first authorship (Ling Zhao, Wei Yang, Yang Chen, Fengjie Huang). Corresponding Author: Zhao-xiang Bian, MD, PhD, Institute of Brain and Gut Research, Chinese Medicine Clinical Study Center, School of Chinese Medicine, 7 Hong Kong Baptist University Road, Kowloon, Hong Kong SAR, China, Wei Jia, PhD, Cancer Biology Program, University of Hawaii Cancer Center, 701 Ilalo Street, Honolulu HI 96813, USA, Xiao-dong Fang, PhD, Bioinformatics Group, The Second Affiliated Hospital of Guangzhou University of Chinese Medicine, 111 Dade road, 510120, Guangzhou, China.

## Abstract

**Objective:** An excess of fecal bile acids (BAs) is thought to be one of the mechanisms for diarrhea-predominant irritable bowel syndrome (IBS-D). However, the factors causing excessive BA excretion remains unclear. Given the importance of gut microbiota in BA metabolism, we hypothesized that gut dysbiosis might contribute to excessive BA excretion in IBS-D.

**Design:** Metabolomic and metagenomic analyses were performed of specimens from 290 IBS-D patients and 89 healthy volunteers. By transplanting human microbiota and manipulating specific microbiome species in mice, the effects of microbiota on host BA metabolism were assessed at metabolic, genetic and protein levels. Effects of individual and mixed BAs on enterohepatic feedback pathways were also tested *in vitro* and *in vivo*.

**Results:** Total fecal BAs were excessively excreted in 24.5% of IBS-D patients. Their fecal metagenomes showed increased abundances of *Clostridia* and BA-transforming genes (*hdhA* and *bais*). The increases of *Clostridia* bacteria (e.g. *C. scindens*) were positively associated with the levels of fecal BAs and serum 7α-hydroxy-4-cholesten-3-one (C4), while being negatively correlated with serum fibroblast growth factor 19 (FGF19). Both *Clostridia*-rich human microbiota and *C. scindens* enhanced levels of serum C4 and hepatic conjugated BAs in mice recipients and reduced ileal FGF19 expression. Inhibition of *Clostridium* species by vancomycin yielded opposite findings. *Clostridia*-derived BAs (e.g. conjugated and free ursodeoxycholic acid) significantly suppressed intestinal FGF19 expression.

**Conclusion:** The *Clostridia*-rich microbiota contributes to excessive BA excretion in IBS-D patients. This study provided the basis for more precise clinical diagnosis and management for IBS-D.

## INTRODUCTION

Irritable bowel syndrome (IBS) characterized by irregular bowel habits affects 10% of the worldwide population,^1^ of which diarrhea-predominant IBS (IBS-D) shows a higher prevalence in Western countries and Asia.^2–4^ IBS imposes a high economic cost on patients and the government.^5^ The efficacy of pharmacologic agents for IBS is variable among patients,^6 7^ which may be the result of differing etiologies between IBS individuals, as well as imperfect understanding of IBS pathogenesis.

As one important pathogenic factor,^8^ fecal bile acids (BAs) have been shown to be excessively excreted in 12% to 43% of IBS-D patients, but less often observed in other IBS subtypes.^9–12^ The increase of total fecal BAs shows close correlations with IBS-D symptoms, such as abdominal pain and accelerated colonic transit time.^13 14^ Animal studies have shown that infusion of BAs, such as deoxycholic acid (DCA), can induce accelerated colonic motility and visceral hypersensitivity.^15 16^ In contrast, attenuation of BA excretion by administration of sequestering agents partially alleviates the symptoms of IBS-D patients.^17 18^ However, the mechanism underlying excessive fecal BA excretion in IBS-D remains unclear.

Generally, primary BAs are synthesized by the rate-limiting enzyme cholesterol 7α-hydroxylase (CYP7A1) and then conjugated with amino acids in the liver.^19^ Most conjugated BAs released into the intestine are reabsorbed from the distal ileum and transported into the portal vein for circulation back to the liver, a process known as enterohepatic circulation.^19^ A small amount of BAs escaping enteric absorption is ultimately excreted in the stool daily, and they can be replenished by hepatic *de novo* synthesis.^20^ The rate of hepatic BA synthesis is reversibly controlled by Farnesoid X receptor (FXR)-mediated feedback mechanisms in the liver and ileum.^21^ Moreover, gut microbiota, as an indispensable participant in BA metabolism, is responsible for conversion of primary BAs into secondary BAs in the GI tract.^20^ And they also control host BA synthesis by affecting intestinal FXR-mediated feedback signaling.^22^

Previous studies have shown the metabolic and genetic disturbances related to the enterohepatic circulation in IBS-D. For example, serum 7α-hydroxy-4-cholesten-3-one (C4), an intermediate product of CYP7A1 that reflects the BA synthetic rate, were significantly increased in IBS-D patients.^23^ Serum fibroblast growth factor 19 (FGF19), an intestine-released hormone that negatively controls BA synthesis, was decreased in patients with functional diarrhea.^24^ Moreover, varied genotypes of FGF19 receptor complex were associated with total fecal BA levels in IBS-D subjects.^11 12^ These phenomena suggest there is a link between excessive BA excretion in IBS-D and enhanced BA synthesis or impaired feedback control of BA synthesis.

Altered gut microbiota has been reported in the IBS-D population,^25 26^ however, its link with excessive BA excretion has received little attention. Given the important role of gut microbiota in BA synthetic regulation and the appearance of gut dysbiosis, we hypothesized that gut dysbiosis may contribute to the excessive BA excretion in IBS-D. To investigate this hypothesis, BA-related metabolomic and metagenomic analyses were performed of an IBS-D cohort to identify the association of gut microbiota with BA disturbance. Further, whether and how gut microbiota affect BA excretion in IBS-D subjects were investigated through a series of animal and cell experiments.

## MATERIALS AND METHODS

### Subject recruitment and sample collection

A total of 345 IBS-D adults meeting Rome IV criteria were prospectively recruited at two Chinese medicine clinics affiliated with the School of Chinese Medicine, Hong Kong Baptist University (HKBU). Based on an approximately 30% of the pooled prevalence of the IBS-D population,^10–12 27^ 91 healthy subjects as controls (HC) were also invited. This study was approved by the Ethics Committee on the Use of Human & Animal Subjects in Teaching & Research of HKBU (Approval no. HASC/15-16/0300 and HASC/16-17/0027). Written consent was obtained from each subject prior to sample collection. All subjects were instructed to provide fasting blood and morning first feces, they were required to stop using antibiotics, probiotics, prebiotics and other microbiota-related supplements at least one month before sampling. Human specimens (serum and stools) were transported to the laboratory using dry ice and were frozen at −80 °C. Details of patients’ diagnoses, subject recruitment and sampling are described in **Supplementary methods.**

### Fecal metagenomic sequencing and data processing

Microbial DNA was extracted from stool samples (200 mg) by the phenol/chloroform/isoamyl alcohol method.^28^ Fecal metagenomes were sequenced in high quality fecal DNA samples. The DNA library was prepared using a TruSeq DNA HT Sample Prep Kit and sequenced by the Illumina Hiseq 2000 at BGI-Shenzhen. Removing low quality bases and human genome left 83.59% of the high-quality sequences; these were mapped using the published gene catalog database of the human gut microbiome.^29^ Microbial diversity was determined and taxa were identified as described previously.^30 31^ Functional orthologs (KOs) were predicted against the KEGG gene database (v79) by BLASTP with the highest scoring annotated hits. The relative abundances of phyla, genera, species and KOs were calculated from the relative abundances of their respective genes.

### Measurement of C4 and FGF19 in serum samples

Concentrations of FGF19 in human serum samples were tested using a commercial Human Fibroblast Growth Factor 19 Assay Kit (Thermo scientific, Waltham, MA, USA). The level of serum C4 was quantified by a mass spectrometry-based method developed by our group.^32^

### Quantification of BAs in specimens of human beings and rodents

BA metabolites were individually extracted and quantified from human samples (serum and stools) and rodents’ specimens (liver and ileal contents) as previously described.^32–35^ With mass spectrometry as the platform, a single 26-min acquisition method with positive/negative ion switching under the multiple reaction monitoring (MRM) mode was used for simultaneous quantification of 36 BA individuals and C4. The chemical usage and detail procedures are described in **Supplementary methods**. The tested BAs with their specific MRM transitions and MS/MS parameters are shown in **Table S1**. Total serum or fecal BA levels of included subjects and animals were obtained by accumulation of all testing BAs.

### Animals

Male C57BL/J mice were purchased from the Laboratory Animal Services Centre of The Chinese University of Hong Kong, Hong Kong. Animals were housed in rooms maintained on a 12-h light/dark cycle with free access to a standard rodent diet and water. Performance of animal experiments was in accordance with the Animals Ordinance guidelines, Department of Health in Hong Kong.

### Transplantation of human microbiota in pseudo germ-free mice

The pseudo germ-free (GF) model was established in eighteen mice (n=6/group) by antibiotic cocktail (ABX) prior to fecal microbial transplantation (FMT).^36^ Human microbiota suspensions (50mg/mL) were separately prepared from stool samples of HC and IBS-D donors (n=11-12/group). The demographics and BA-related quantitative traits of donors were shown in **Table S2**. ABX-pretreated mice were daily gavaged with 200 μL of fecal suspension for five consecutive days. One week after FMT, GI motility and fecal consistency were measured using our previous protocol^37^. Fecal samples were collected at baseline, before and after FMT for monitoring bacterial density. Finally, after anesthesia, cecal contents were collected for testing BA-transforming bacterial abundances and activities. Other specimens (liver, ileum, and ileal contents) were collected to analyze BAs and/or BA-related genes and proteins. Measurements of genes, proteins and BA transforming activity of donors’ and recipients’ microbiota are shown in **Supplementary methods**.

### Manipulation of *Clostridium* species in conventional mice

Eighteen mice (n=6/group) were used for this experiment. A typical BA-transforming *Clostridium* strain (*C. scindens* ATCC 35704) was purchased from the American Type Culture Collection and was identified by its 16s ribosomal sequence. The strain was cultured using Brain Heart Infusion (BHI) broth under anaerobic conditions (Bactron300 Anaerobic Chamber Glovebox, Shel Lab Inc., USA). After confirmation of its BA converting activity *in vitro*, *C. scindens* PBS suspension (10^8^ CFU/mL) was daily gavaged to one group of mice last for 7 days. With antibacterial activity against *Clostridium*,^38^ vancomycin (0.1 mg/mL) was administrated to another group of mice for inhibition of *Clostridium* species. Vehicle controls were gavaged with PBS. BA-related bowel and metabolic phenotypes were done after stopping bacterial manipulations as mentioned above in the FMT experiment.

### BA intervention in mice

Forty mice were divided into 5 groups of 8 mice. Each of the four experimental groups was gavaged with 50mg/kg body weight of either TCA, TCDCA, TUDCA individuals or their mixture (T-BAs) for 8 weeks. Control mice in the fifth group were treated with saline. The intervention dose and duration were as in a previous study.^39^ After anesthesia, tissues of liver and ileum were collected for analyzing genes of FXR, FGF19, small heterodimer partner (SHP) and BA synthases (CYPs) by real-time PCR.

### Culture and treatment of human hepatocytes and enterocytes

Human liver cell line L02 and intestinal cell line NCI-H716 purchased from ATCC were cultured with Dulbecco’s Modified Eagle’s Medium (DMEM) and Roswell Park Memorial Institute (RPMI) 1640 Medium with 10% fetal bovine serum (FBS), respectively. L02 cells were treated with vancomycin (10 µM to 400 µM) and *C. scindens* (live and heat-killed) for 24 hours to test their effects on CYP7A1 expression. Based on the EC_50_ range of BAs for FXR activation,^40^ hepatocytes and enterocytes were separately treated with 50μM of BA individuals or mixtures for 24 hours to analyze gene or protein expressions of FXR, FGF19, SHP and CYPs.

### Statistical analysis

Software packages R 3.4.3 (https://www.r-project.org/) was used to perform the Shapiro-Wilk’s test for judging the distribution of human total fecal BAs. The relationships of total fecal BAs with other biochemical and bacterial characteristics were analyzed by Spearman’s correlation based on Prism 7 (GraphPad Software, La Jolla, CA). Differential taxa and BA-transforming genomes were analyzed with the Benjamin-Hochberg method. Variations of clinical quantitative traits, metabolites and genes were analyzed by the nonparametric Kruskal-Wallis tests for comparison of multiple groups while the Mann-Whitney test was employed for comparison of two groups. The significance difference for all of tests was set to *p*< 0.05.

## RESULTS

### Higher levels of BA synthesis in the subgroup of IBS-D patients with excessive BA excretion

A total of 345 IBS-D adults meeting ROME IV criteria were recruited, of which 290 patients completed all biochemical tests and consented to voluntarily provide biospecimens for the current study. Meanwhile, 91 age- and gender-matched HC subjects were recruited, of whom 89 provided biospecimens. As shown in Table 1, the IBS-D cohort displayed a significant increase in defecation frequency and total fecal BAs whereas decreased fecal consistency compared with HC subjects. The level of total fecal BA excretion showed a skewed distribution (*p*<0.05 by the Shapiro-Wilk’s test) in IBS-D and HC group (Figure 1A).

**Figure 1.**
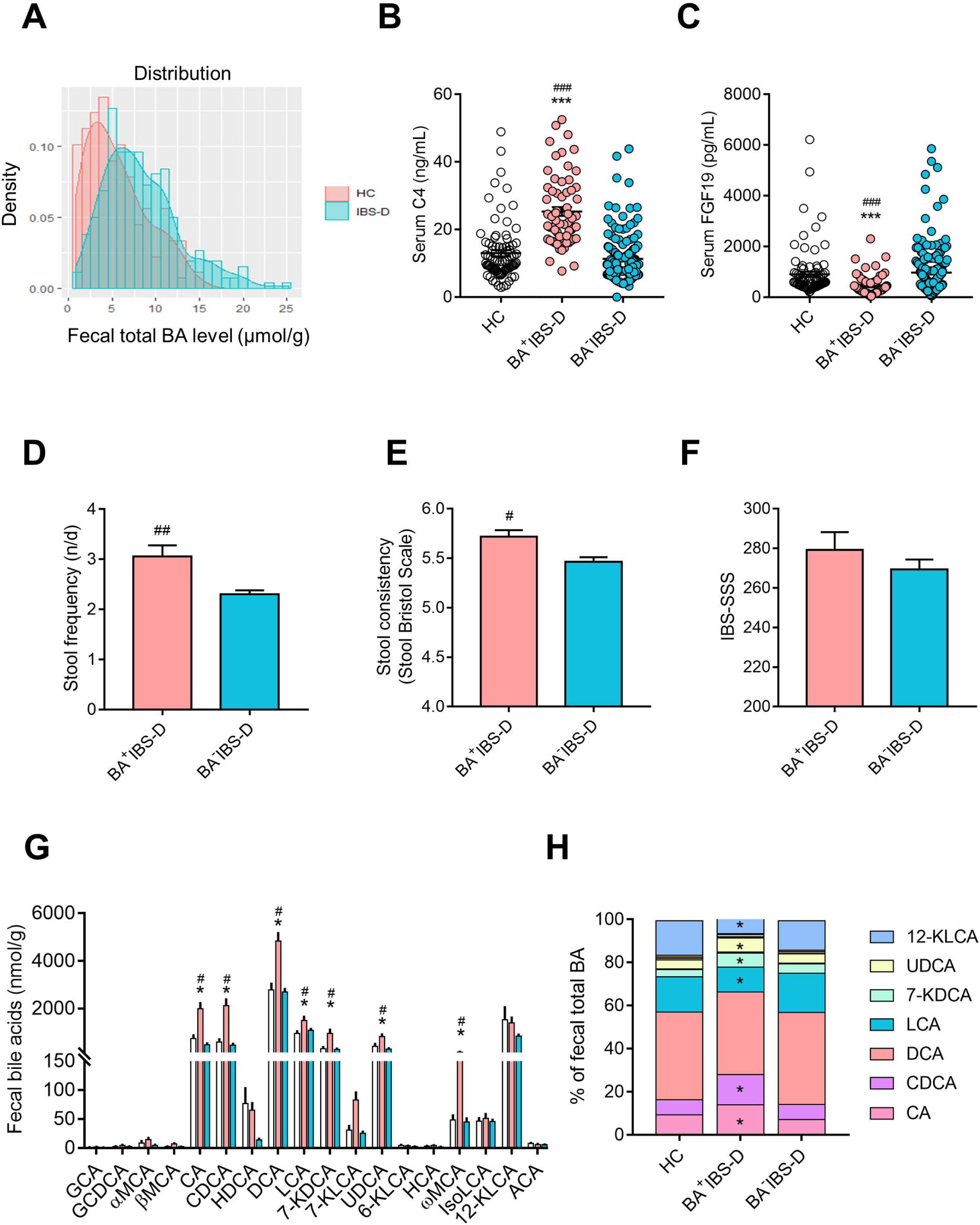
Alteration of fecal BA profiles and serum BA synthetic indicators in IBS-D patients. (A) The histogram of the distribution of total fecal BA level in healthy controls (n=89) and IBS-D patients (n=290). Based on the 90^th^ percentile of healthy total fecal BA level, a quarter of IBS-D patients (n=71) with excessive BA excretion were grouped as BA^+^IBS-D and other patients (n=219) were classified as BA^-^IBS-D. (B, C) Concentrations of serum 7α-Hydroxy-4-cholesten-3-one (C4) and fibroblast growth factor 19 (FGF19). (D-F) The severity of bowel symptoms between IBS-D subgroups assessed by defecation frequency, Stool Bristol Scale and IBS Severity Scoring System (IBS-SSS). (G) Absolute contents of fecal dominant BAs. (H) Proportions of fecal dominant BAs. Only BAs with over 1% of total BA pool were shown in the legend. Difference in phenotypic scores between IBS-D subgroups were analyzed by the Mann-Whitney test, and BA-related indices were evaluated among three groups by the Kruskal-Wallis test. Statistical significance is expressed by *, *p*<0.05; ***, *p*<0.005 compared with the HC group; #, *p*<0.05; ##, *p*<0.01; ###, *p*<0.005 compared with the BA^-^IBS-D group.

**Table 1.**
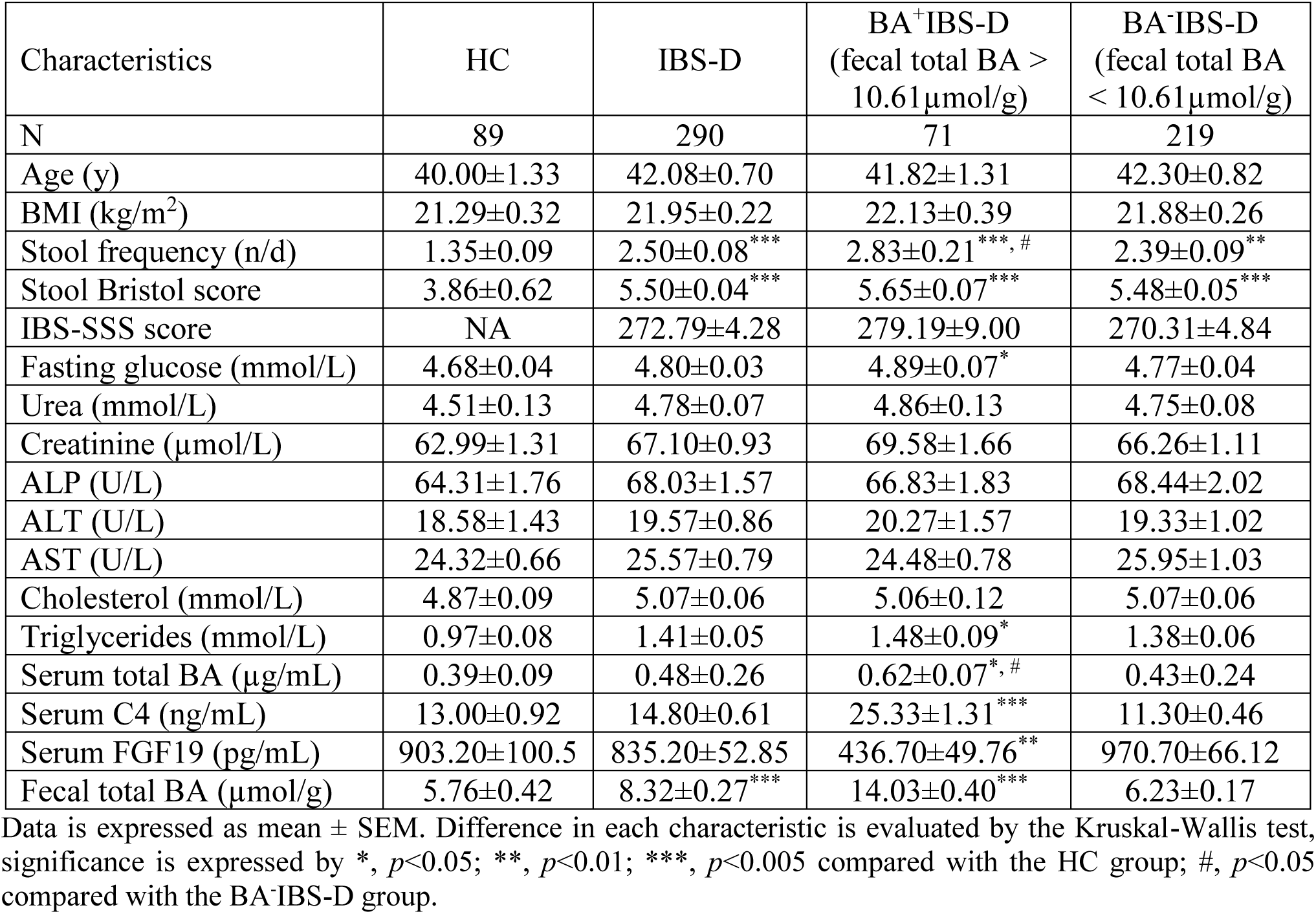
The demographics and clinical characteristics of IBS-D patients based on total fecal BA excretion.

A percentage of 24.5% of IBS-D patients (71 of 290) was found to have an excess of total BA excretion in feces (≥10.61µmol/g) by the 90^th^ percentile cut-off value as determined from the HC group; these were classified as BA^+^IBS-D. Remaining patients with normal fecal BA excretion (<10.61µmol/g) were grouped as BA^-^IBS-D. Comparing with HC and BA^-^IBS-D groups, BA^+^IBS-D patients also exhibited increased C4 and decreased FGF19 in serum, as well as increased severity of diarrheal symptoms (Table 1; Figure 1B-F). Correlation analysis revealed that, in the BA^+^IBS-D group, total fecal BA levels were positively associated with serum C4 levels and scores of diarrheal symptoms (Bristol Stool Scale and defecation frequency), while being inversely correlated with serum FGF19 levels (**Table S3**). These results demonstrated that enhanced BA synthesis was occurring in IBS-D patients, accompanied by excessive BA excretion and increased severity of diarrheal symptoms.

Alteration of individual BAs was also specifically observed in the serum and feces of BA^+^IBS-D patients, but no difference was detected in BA^-^IBS-D patients. Serum BA profiles revealed that conjugated BAs and free primary BAs largely dominated the total serum BA pool in this cohort. Particularly, glycochenodeoxycholic acid (GCDCA), glycoursodeoxycholic acid (GUDCA) and chenodeoxycholic acid (CDCA) showed significant elevation in both absolute amounts and relative proportions in BA^+^IBS-D patients relative to the HC group (Figure S1). In addition, BA^+^IBS-D patients had an increased absolute level of ursodeoxycholic acid (UDCA) and a reduced relative proportion of glycohyodeoxycholic acid (GHDCA) in serum. Moreover, the fecal BA pool of recruits was largely comprised of free BA metabolites as previously described,^41^ of which cholic acid (CA), CDCA, DCA, lithocholic acid (LCA), 7-ketodeoxycholic acid (7-KDCA), UDCA and ω-muricholic acid (ωMCA) showed significant increases in their absolute amounts in the BA^+^IBS-D group relative to the HC group (Figure 1G). Meanwhile, proportions of CA, CDCA, UDCA and 7-KDCA were also showed increased in total fecal BAs whereas proportions of LCA and 12-KLCA were decreased (Figure 1H). But there was no difference in fecal BA composition between the BA^-^IBS-D and HC groups. Results of serum and fecal BA profiles revealed altered composition of microbiota-derived BAs in BA^+^IBS-D patients, indicating there might be an abnormality in BA-transforming microbiota.

### Association of a *Clostridia*-rich microbiota with increased BA synthesis and excretion in BA^+^IBS-D patients

Fecal metagenomic data was successfully obtained from 84 HC subjects, 70 BA^+^IBS-D patients and 207 BA^-^IBS-D patients. An average 6.04 GB of high-quality sequencing reads was obtained from each sample, and an average 63.7% of reads per sample were successfully mapped (**Table S4**). In comparison to the HC and BA^-^IBS-D groups, BA^+^IBS-D patients’ fecal microbial communities showed a higher Bray-Curtis dissimilarity index with no difference in total gene count and Shannon index (Figure 2A **& S2A-B**). These results indicate unchanged microbial richness within all included subjects, but with a larger instability of the enteric ecosystem in the BA^+^IBS-D subgroup.

**Figure 2.**
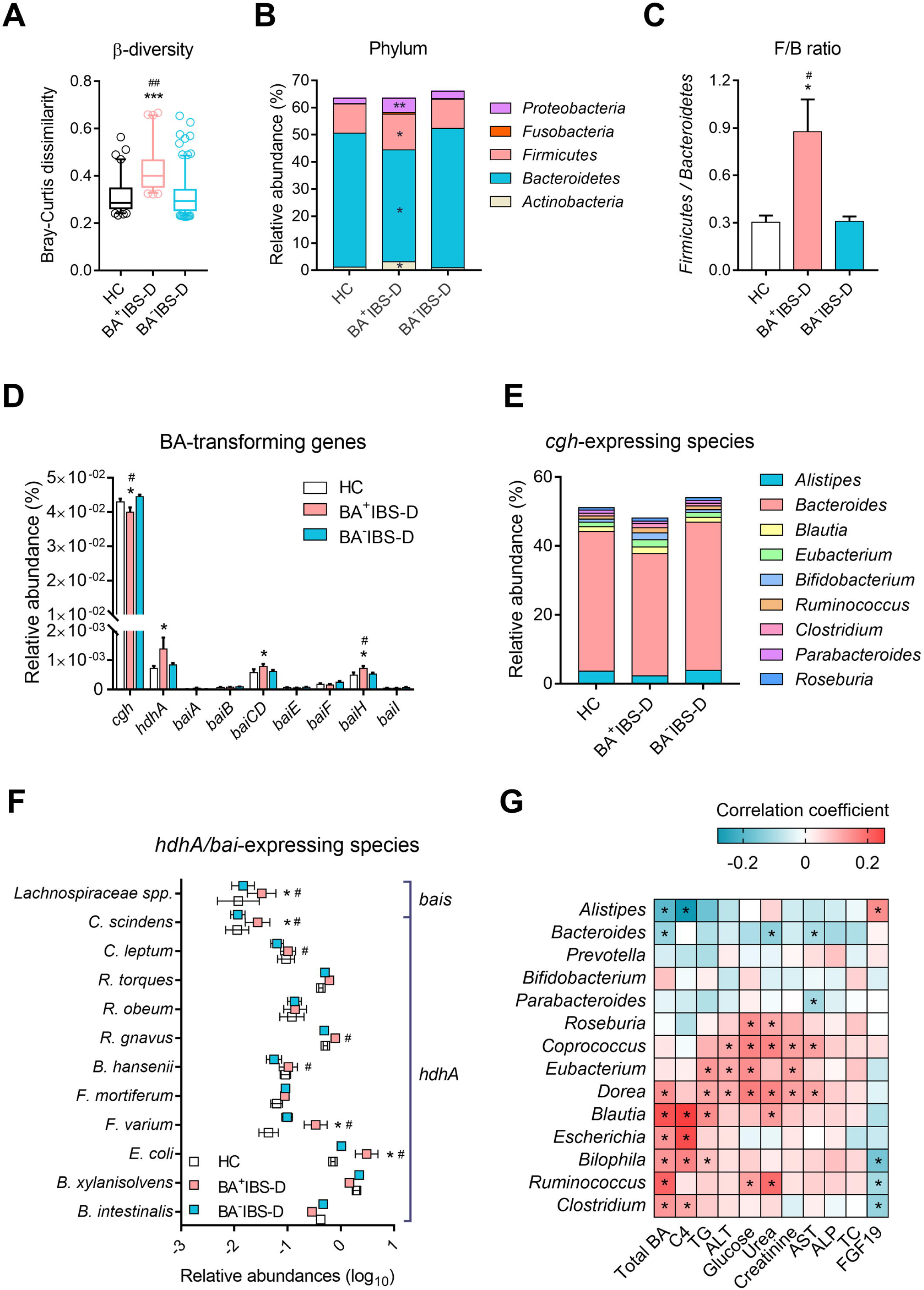
The association between the *Clostridia*-rich microbiota and the levels of BA synthesis and excretion in IBS-D cohort. (A) Microbial β-diversities measured by Bray-Curtis dissimilarity. (B, C) The relative abundances of dominant phyla and the ratio of *Firmicutes* to *Bacteroidetes* (F/B). (D-F) The relative abundances of BA-transforming genomes and bacteria. (G) Spearman’s correlation between bacterial abundances and biochemical indexes in the IBS-D cohort. Metagenomic dataset was obtained from 277 IBS-D and 84 HC fecal samples. Differential taxa and genes among three groups were analyzed with the Benjamin-Hochberg method, statistical significance is expressed by *, *p*<0.05; ***, *p*<0.005 compared with the HC group; #, *p*<0.05; ##, *p*<0.01 compared with the BA^-^IBS-D group. Statistical significance for Spearman’s correlation is set as *, *p* < 0.05.

Principal coordinate analysis revealed an overlapping of fecal microbial communities among IBS-D and HC subjects (**Figure S2C**), however, a distinct microbial profile was shown in BA^+^IBS-D patients in comparison with either HC subjects or BA^-^IBS-D at different taxonomic levels (**Table S5-7**). Specifically, the phyla *Firmicutes*, *Actinobacteria*, *Fusobacteria* and *Proteobacteria* showed increased relative abundances while *Bacteroidetes* exhibited a decreased relative abundance in BA^+^IBS-D patients (Figure 2B). The ratio of *Firmicutes* to *Bacteroidetes* (F/B) was significantly elevated similarly (Figure 2C). At genus level, several *Clostridial* bacteria, including *Ruminococcus*, *Clostridium*, *Eubacterium* and *Dorea*, showed increased abundances in BA^+^IBS-D fecal microbiota (**Figure S2D**). *Bifidobacterium*, *Escherichia* and *Bilophila* were also more abundant. However, the abundances of *Alistipes* and *Bacteroides* were reduced.

The alteration in the bacterial composition of the BA^+^IBS-D group was also associated with variation in BA-transforming genomes (Figure 2D**, Table S8**). Reduced abundances of *Alistipes* and *Bacteroides* were mainly associated with a decreased abundance of the *cgh* gene (Figure 2E), which encodes the BA deconjugating enzyme choloylglycine hydrolase.^41^ However, an elevated abundance of the *hdhA* gene, encoding 7α-hydroxysteroid dehydrogenase (7α-HSDH),^41^ was attributed to increases of *Escherichia*, *Fusobacterium*, *Blautia*, *Ruminococcus* and *Clostridium* species (Figure 2F). Moreover, *Clostridium scindens* (*C. scindens*) and an unclassified *Lachnospiraceae* species largely contributed to higher abundances of genes *baiCD* and *baiH* (Figure 2F), that are known to encode 7-dehydroxylases.^41^

Correlation analysis revealed that abundances of *Clostridia* bacteria (including *C. scindens*) showed positive relationships with levels of total fecal BA and serum C4, but inverse correlations with the serum FGF19 level (Figure 2G **& S2E**). Metagenomic results identified a specific *Clostridia*-rich microbiota in the BA^+^IBS-D group, with differential genomes for BA deconjugation, C7-isomeric and dehydroxylating actions. Given the importance of gut microbiota in maintenance of host BA metabolism,^19^ the close relationships between bacteria and biochemical indices suggested that the *Clostridia*-rich microbiota may influence IBS-D patients’ BA synthesis and excretion.

### Enhancement of BA synthesis and excretion in pseudo GF mice transplanted with *Clostridia*-rich fecal microbiota

To investigate the effects of the *Clostridia*-rich microbiota on BA metabolism, transplantation of human fecal microbiota was performed in ABX-induced pseudo GF mice (Figure 3A **& S3A**). One week after FMT, similar profiles of fecal BA-transforming bacteria were found between BA^+^IBS-D microbiota recipients and their donors, with a reduced level of *Bacteroidetes* and elevated levels of *Firmicutes*, *Clostridium cluster XIVa* (*C. XIVa*) and *C. scindens* (Figure 3B). Meanwhile, cecal microbiota isolated from BA^+^IBS-D recipients was found to exhibit a significant decrease of deconjugating capability but slight increases of 7-HSDH and 7α-dehydroxylating levels (**Figure S3B**). These *in vitro* transforming results were consistent with what was determined in the fecal microbiota of the human BA^+^IBS-D donors (**Figure S3C**). Microbial results demonstrated that the *Clostridia*-rich microbiota was replicated in the ABX mice receiving human BA^+^IBS-D fecal microbiota.

**Figure 3.**
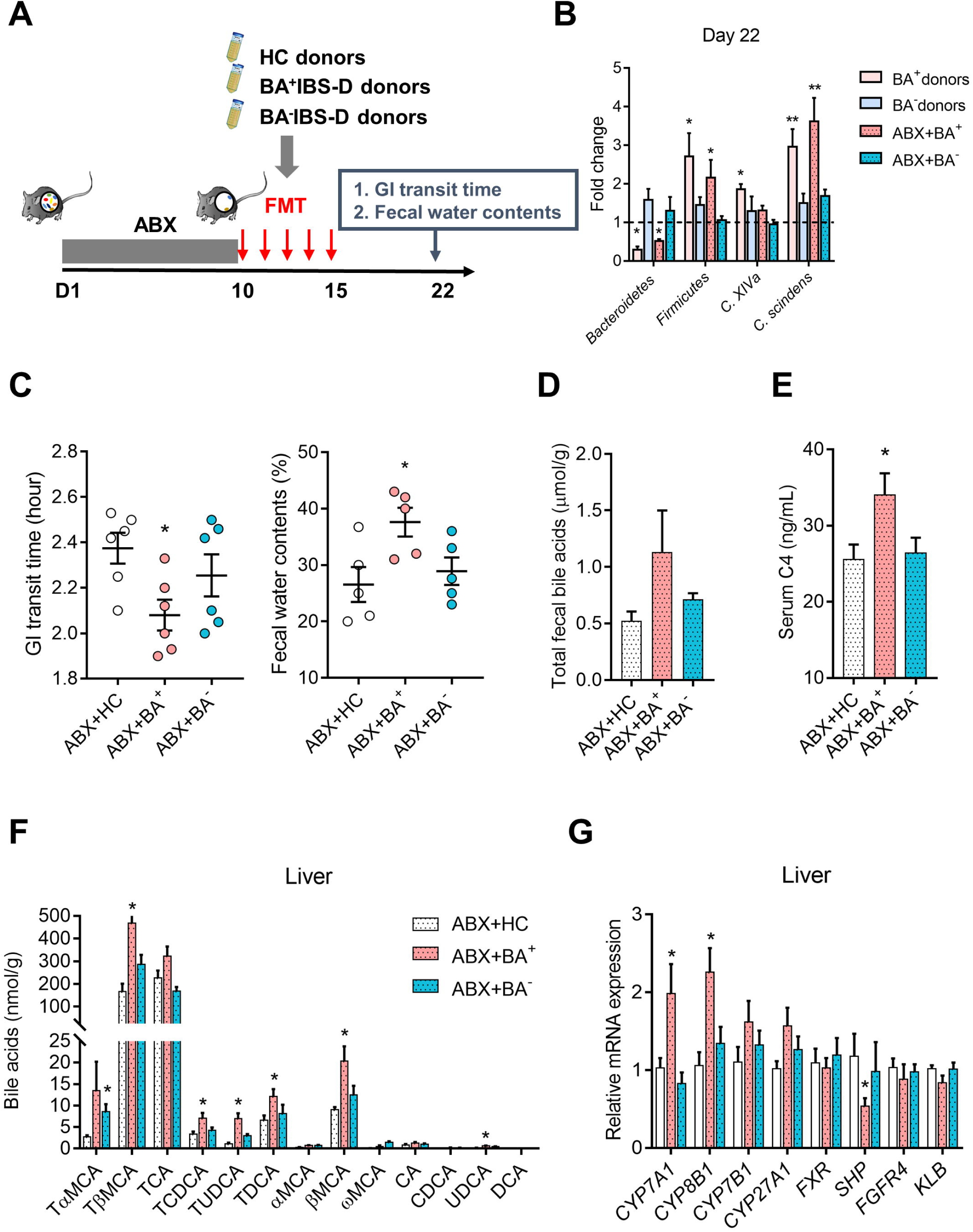
Excessive BA synthesis and excretion in mouse recipients receiving BA^+^IBS-D fecal microbiota. (A) Experimental procedure for fecal microbiota transplantation (FMT) in antibiotic cocktail (ABX)-induced pseudo germ-free mice (n=6/group). Mice that received fecal microbiota of HC donors were grouped as ABX+HC, mice treated with fecal microbiota from BA^+^IBS-D and BA^-^ IBS-D donors were classified as ABX+BA^+^ and ABX+BA^-^, respectively. (B) Relative levels of BA-related bacteria in feces of donors and mouse recipients based on qPCR analysis. (C) The GI transit time and fecal water contents of mouse recipients. (D, E) The levels of total fecal BAs and serum C4 in mouse recipients. (F) Hepatic BA profiles of mouse recipients. (G) Relative gene expression of BA synthetic regulators in the hepatic tissues of mouse recipients. Differential BA-related phenotypes, bacteria, metabolites and genes were assessed with the Kruskal-Wallis test, and statistical significance is expressed by *, *p*<0.05; **, *p*<0.01 compared with the ABX+HC group.

Meanwhile, BA^+^IBS-D microbiota recipients displayed BA-related diarrheal symptoms, including shortened GI transit time and decreased fecal consistency (Figure 3C), that were similar to the symptoms of their donors. Metabolic results found increased amounts of total fecal BAs and serum C4 in the BA^+^IBS-D microbiota recipients, along with elevated taurine-conjugated BA metabolites (TβMCA, TCA, TCDCA and TUDCA) in the liver and ileal lumen (Figure 3D-F **& S3D**). Furthermore, results of hepatic mRNA analysis showed elevated expressions for the BA synthetases, CYP7A1 and CYP8B1, in the BA^+^IBS-D microbiota recipients (Figure 3G), along with reduced SHP mRNA expression. Similarly, the ileal feedback hormone, FGF15, also showed significant reduction in its mRNA expression in BA^+^IBS-D microbiota recipients (**Figure S3E**). Increased expression of hepatic CYP7A1 and decreased expression of ileal FGF15 were also validated at protein level (**Figure S3F**). Additionally, hepatic FGF19 receptor complexes (Fibroblast Growth Factor Receptor 4, FGFR4; Klotho beta, KLB) and ileal BA active transporters (apical sodium bile acid transporter, ASBT; multidrug resistance associated protein 2/3, MRP2/3; and organic solute transporter α/β, OSTα/β) were unchanged among recipient groups. These results suggested that the *Clostridia*-rich microbiota of BA^+^IBS-D donors could enhance BA synthesis, excretion and, induce BA-related diarrhea-like symptoms in mouse recipients, and such microbiota-induced effects might be independent of ileal BA transport and hepatic feedback signal reception.

### Enhancement of hepatic BA synthesis and excretion in mice with colonization of *Clostridium* **species**

To further clarify the effects of *Clostridium* species on BA synthesis and excretion, we manipulated the *Clostridium* species in wild-type mice (Figure 4A). Compared with vehicles, colonization of *C. scindens* (10^8^ CFU/mL) increased the abundance of *C. scindens* in mouse cecal microbiota (Figure 4B). The cecal microbiota of mice with *C. scindens* colonization showed a slight reduction of deconjugating activity and a significant enhancement of 7-HSDH activity compared with that of the PBS group **(Figure S4A**). In contrast, the cecal microbiota of vancomycin-treated mice showed a significant attenuation in the abundances of *Clostridium* species, along with reduced *in vitro* activities of 7-HSDH and 7-dehydroxylases (**Figure S4A & 4B**). However, BA deconjugating capability was remarkably enhanced. Moreover, colonization of *C. scindens* decreased fecal consistency and, slightly accelerated the GI transit in mice relative to vehicles, but inhibition of *Clostridium* by vancomycin significantly attenuated mouse GI movement (Figure 4C).

**Figure 4.**
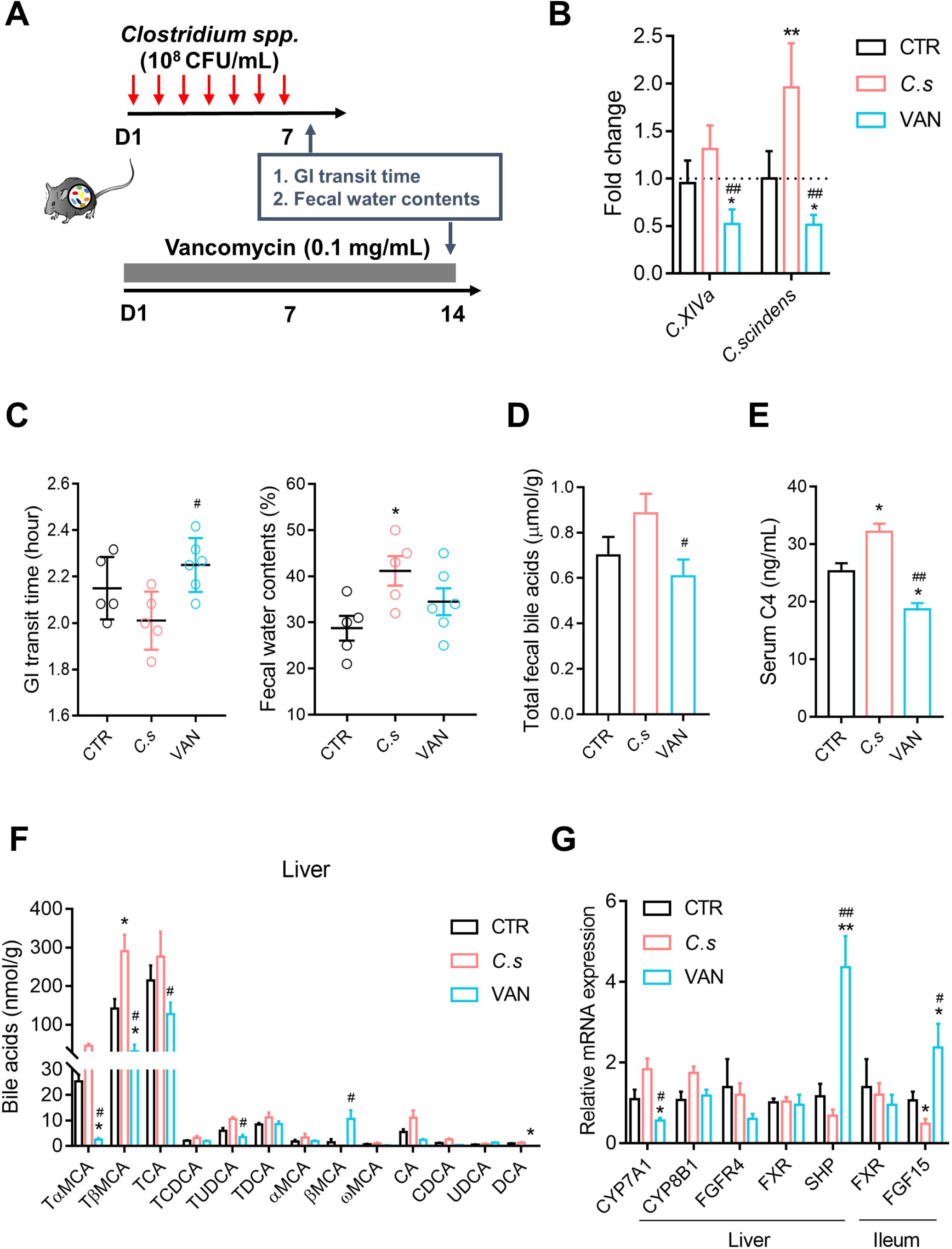
Dysregulation of BA synthesis and excretion in mice with manipulation of *Clostridium* species. (A) Manipulation of *Clostridium* species in wild-type mice (n=5-6/group) through introduction of *C. scindens* strain (*C.s*) or administration of vancomycin (0.1 mg/mL). (B) Relative levels of the fecal *Clostridial* bacteria (*C. XIVa*) and *C. scindens* measured by qPCR analysis. (C) The GI transit time and fecal water contents of *Clostridium*-treated mice. (D, E) The levels of total fecal BAs and serum C4. (F) The BA profile in the mouse liver (G) Relative gene expression of BA synthetic regulators in the hepatic tissues. Differential BA-related phenotypes, bacteria, metabolites and genes were assessed with the Kruskal-Wallis test, and statistical significance is expressed by *, *p*<0.05; **, *p*<0.01 compared with the control group; #, *p*<0.05; ##, *p*<0.01 compared with the *C. s* group.

Metabolic analysis found that amounts of total fecal BAs and serum C4 were significantly increased in *C. scindens-*colonized mice while reduced in *Clostridium*-deficient mice induced by vancomycin (Figure 4D-E). Consistently, taurine-conjugated BAs (TβMCA, TCA and TUDCA) exhibited higher contents in the liver and ileal lumen of *C. scindens-*colonized mice but were significantly reduced in *Clostridium*-deficient mice (Figure 4F **& S4B**). Results of BA synthetic genes revealed that *C. scindens* elevated CYP7A1 mRNA expression in the mouse liver without significance (*p*<0.08). However, vancomycin-treated mice exhibited a significant reduction in the gene expression of hepatic CYP7A1 (Figure 4G). Although FXR gene expression showed no difference within groups with bacterial manipulation, its mediated feedback regulators, hepatic SHP and ileal FGF15, both showed dramatic increases in their gene expression upon vancomycin treatment. Introduction of *C. scindens* significantly attenuated FGF15 gene expression only, which was also verified at protein level (Figure 4G **& S4C**). The FGF15 receptors FGFR4 and KLB showed no difference in expression levels among groups (Figure 4G). These findings demonstrated that introduction of *Clostridium* species led to enhanced BA synthesis and excretion while inducing diarrheal phenotypes in wild-type mice. Inhibition of *Clostridium* species with antibiotics yielded opposite findings. The BA-related changes in the *C. scindens*-colonized mice were also similar to what was detected in BA^+^IBS-D fecal microbiota recipients.

The unchanged expression of CYP7A1 in hepatocytes with treatment of either vancomycin (from 10 to 400 µM) or *C. scindens* strains (live and heat-killed) excluded their direct effects on hepatic BA synthesis (**Figure S4D-E**). Given the inhibitory impact of natural BA products on FXR activation,^42^ we supposed that the excess of BAs (TβMCA, TCA and TCDCA and TUDCA), consistently detected in mice colonized with *Clostridia*-rich microbiota and *C. scindens* may inhibit FXR-mediated feedback signaling, thus enhancing hepatic BA synthesis.

### Inhibitory effects of *Clostridia*-derived BAs on the intestinal negative feedback signalling

Among dominant BAs detected in mice with the *Clostridia*-rich microbiota, TβMCA has been previously shown to antagonize FXR.^22^ We therefore examined the effects of other taurine-conjugated BAs, TCA, TCDCA, TUDCA and their mixture (T-BAs) on hepatic and intestinal FXR feedback pathways *in vitro* and *in vivo*.

The doses of BAs (50 µM) used in cell experiments were based on a published EC _50_ range for FXR ligands.^40^ The experiments with NCI-H716 cells revealed that TUDCA and T-BAs slightly attenuated FXR expression and significantly reduced FGF19 expression in enterocytes (Figure 5A). In hepatocytes (L02 cells), we found that T-BAs induced a significant reduction in FXR expression (**Figure S5A**). Also, TUDCA and T-BAs can reduce SHP expression, but only the former significantly elevated expressions of CYP7A1 in hepatocytes. We further confirmed these effects *in vivo* using an 8-week BA intervention (50mg/kg/d) in mice. Ileal FXR gene expression was significantly reduced by TUDCA and slightly attenuated by TCDCA and T-BA. The three treatments significantly decreased ileal FGF15 expressions in mice but elevated hepatic CYP7A1 expressions (Figure 5B **& S5B**). In mouse liver, taurine-conjugated BAs had no effects on FXR expression, and TUDCA increased SHP expression (**Figure S5B**). These results showed that mouse *Clostridia*-derived BAs (TUDCA and T-BAs) can attenuate intestinal FXR/FGF15 signaling, but their effects on hepatic FXR/SHP signaling were controversial.

**Figure 5.**
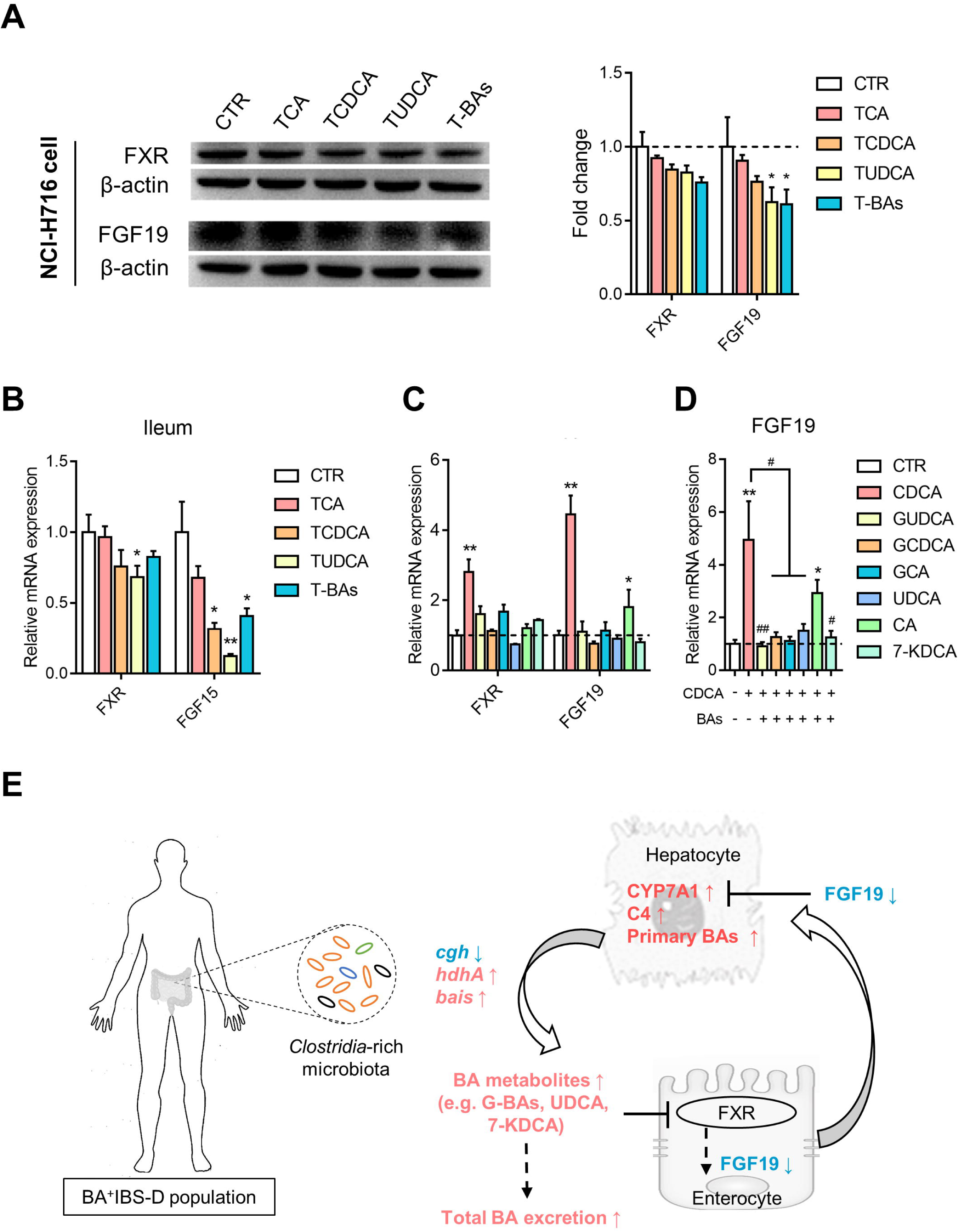
The inhibitory effects of *Clostridia*-derived BAs on intestinal FXR feedback signaling. (A) Expressions of FXR and FGF19 in enterocytes treated with BA taurine-conjugates that were derived from the mouse *Clostridia*-rich microbiota. T-BAs represents a combination of TCA, TCDCA and TUDCA. (B) Gene expression of ileal FXR and FGF19 in mice with intervention of taurine-conjugated BAs (n=8/group). (C) Gene expression of FXR and FGF19 in enterocytes treated with individual BAs that were derived from the human *Clostridia*-rich microbiota. (D) Gene expression of FGF19 in enterocytes treated with combination of FXR agonist CDCA and each *Clostridia*-derived BAs. (E) Schematic diagram of a potential mechanism by how the *Clostridia*-rich microbiota contribute to excessive BA synthesis and excretion in IBS-D. Differential proteins or genes were analyzed with the Kruskal-Wallis test, and statistical significance is expressed by *, *p*<0.05; **, *p*<0.01 compared with the control group; #, *p*<0.05; ##, *p*<0.01 compared with the CDCA group.

To further clarify whether BAs derived from human *Clostridia*-rich microbiota also affect intestinal FXR/FGF19 signaling, we examined FXR and FGF19 gene expressions in human enterocytes with different interventions. CDCA, as the most potent agonist of FXR,^43^ dramatically elevated FXR and FGF19 expressions in enterocytes (Figure 5C). CA also increased FGF19 expression. Although single treatment of other BAs (GCDCA, GUDCA, GCA, UDCA and 7-KDCA) exhibited no effects on FXR and FGF19 expressions, these BAs can efficiently antagonize CDCA-induced FXR activation in the enterocytes (Figure 5C-D). Taken together, these results revealed that specific composition of BAs derived from either mouse or human *Clostridia*-rich microbiota can inhibit intestinal FXR/FGF19 signaling.

## DISCUSSION

In this study, we found that a *Clostridia*-rich microbiota was positively associated with increased BA synthesis and excretion in an IBS-D cohort. The causal relationships between the *Clostridia*-rich microbiota and enhanced host BA synthesis and excretion was further confirmed in animals subjected to different bacterial manipulations. Mechanistic experiments both *in vitro* and *in vivo* revealed that *Clostridia*-derived BAs had an attenuated impact on intestinal BA feedback inhibition.

Excessive BA excretion is one of the most characteristic clinical manifestations for IBS-D patients with primary bile acid diarrhea (BAD). In accordance to existing pathogenic understandings, fecal BAs, or combination with serum BA synthetic markers, has been used as auxiliary indexes for diagnosis of primary BAD in clinical practices.^44^ Our findings among this cohort study showed that 24.5% of IBS had excessive BA excretion, thus supporting the idea that fecal BAs can be used as an auxiliary index for stratifying IBS-D populations. Further, our study also showed that the abundances of *Clostridia* bacteria (such as *C. scindens*) directly contributed to the increase of fecal BAs. Such close relationships suggest that not just excretory BAs but also their related bacteria could be another auxiliary biomarkers for distinguishing primary BAD from IBS-D.

In fact, the microbiota-based concept for subtyping IBS has been previously proposed by Jeffery, *et al.*^45^ Another study also has classified different enterotypes within the IBS category, linked with severity of symptoms, but authors pointed out the difficulty in stratifying IBS patients from healthy subjects when using only classic ecologic approaches^46^. Scrutinizing the linkage of microbial signature with the IBS subtype, varied results have been reported across studies^26^, a reduced microbial diversity and increased F/B ratio were more frequently found.^45 47 48^ Our study did find an increased F/B ratio in this IBS-D cohort, but with no difference in the microbial richness. Notably, proportions of taxa were significanlty altered shown in BA^+^IBS-D patients, not in all IBS-D population, illustrating abnormal BA excretion closely related to differentiation of gut microbiota among the IBS-D population. Microbial structures deserve to be explored to see whether it can differentiate the IBS population, considering the gut microbiota always take effects through its metabolites, accurate characterization of IBS microbial signatures should include their metabolic functions. A specific IBS-D microbial signature combined with its metabolic features, *Clostridia*-rich/BA-excessive, may also benefit the precision of the existing symptom-based diagnosis criteria of IBS. More importantly, such integrating signature can provide a pathogenetic explanation underlying excessive BA excretion in IBS-D.

Previous studies with IBS-D or functional diarrhea have revealed increased C4 and decreased FGF19 levels in sera of patients,^12 17 23 27 49^ leading to a pathogenic hypothesis that excessive BA excretion is due to enhanced BA synthesis caused by deficiency of FGF19 release.^50^ We also found that elevated C4 and reduced FGF19 levels in sera of BA^+^IBS-D patients were significantly associated with total fecal BAs. Several studies have reported unchanged genotype and mRNA expression of ileal transporter ASBT in IBS-D patients, together with similar BA transport across the ileal brush border membrane in diarrheal patients and controls.^11 51 52^ These findings excluded impact of the ileal active transport on enhanced BA synthesis and excretion. Despite lacking tissue-level data, our results of higher amounts of total and glycine-conjugated primary BAs in sera of BA^+^IBS-D patients suggest that the decreased FGF19 level is unrelated to ileal BA transport as well. Further, our study with bacterial manipulations in mice demonstrated that excessive BA synthesis and excretion was primarily contributed by *Clostridia*-rich microbiota.

Altered gut microbiota has been shown to modulate hepatic BA synthesis depending on intestinal FXR/FGF19 feedback signaling.^53^ Sayin, *et al.* reported that increase of TMCAs caused by weakened bacterial deconjugation in GF mice can attenuate intestinal FXR signaling, thus enhancing hepatic BA synthesis.^22^ In our work, inhibitory effects on intestinal FXR/FGF19 signaling were observed in mice colonized with *Clostridia*-rich microbiota and *Clostridium* species. In contrast, mice with elimination of *Clostridium* species presented elevated intestinal FXR/FGF19 signaling. These results indicate the FXR/FGF19 signaling is indeed affected by *Clostridium* species. How do the *Clostridium* species modulate FXR/FGF19 signaling? Our data, in line with previous studies, indicate an indispensable role of intestinal FXR/FGF19 signaling in gut microbiota-mediated hepatic BA synthesis, in both physiological and pathophysiological conditions. The fact that BA metabolites are natural ligands for FXR argues strongly for the conclusion that the effect of *Clostridia*-rich microbiota on intestinal FXR/FGF19 signaling is mediated by BA products derived from specific microbial biotransformation.

BA biotransformation of IBS gut microbiota has been put less attention, with only one study showing a lower deconjugating activity of fecal bacteria isolated from IBS patients relative to controls.^13^ Consistently, we found that the *Clostridia*-rich microbiota in human and mice showed a reduced *in vitro* deconjugating capability, along with higher levels of conjugated BAs in BA^+^IBS-D patients (GCDCA, GCA and GUDCA) and colonized mice (TβMCA, TCA, TCDCA and TUDCA). These findings are in line with low abundances of *cgh* genomes in BA^+^IBS-D microbiota. Moreover, the *Clostridia*-rich microbiota with enrichment of *hdhA* and *bai* genomes exhibited enhanced *in vitro* activities of 7-HSDH and 7-dehydroxlases. Elevated amounts of secondary BAs GUDCA, UDCA, DCA and LCA identified from the BA profile of BA^+^IBS-D patients also supported it. Interestingly, except the capacity of 7α-dehydroxylation for production of DCA and LCA, *C. scindens* was recently discovered to oxidize other hydroxyl groups and reduce ketone groups in primary and secondary bile acids *in vitro*, thus generating keto-BAs and isoBAs.^54^ It suggests that the increase of 7-KDCA and 7-KLCA contents in fecal BA pool of BA^+^IBS-D patients is also contributed by enrichment of 7α-dehydroxylating *Clostridium* species. These observations illustrated specific compositions of secondary BAs with *Clostridia*-rich microbiota in the host. Further, these secondary BAs derived from human and mouse *Clostridia*-rich microbiota both had substantial inhibition in intestinal FXR/FGF19 signaling. Particularly, TUDCA which is dominant in the BA pool was found to reduce FXR and FGF19 expression *in vitro* and *in vivo*. GUDCA, UDCA, 7-KDCA showing higher proportions in the BA pool of BA^+^IBS-D patients suppressed CDCA-activated FXR/FGF19 signaling *in vitro*. Recently, a study with morbidly obese NAFLD patients also reported the suppressive effects of UDCA on circulating FGF19 and FXR activation.^55^ Taking together, our combination of genomic and metabolomic results of microbial BA-transforming actions functionally characterized IBS-D gut microbiota, also gave an explanation why deficient FGF19 release appears in the BA^+^IBS-D group, but not other IBS-D patients.

Moreover, the FXR/SHP signaling in liver, known to negatively regulate gene transcription of hepatic BA synthetase,^56^ was also attenuated in mice colonized with *Clostridia*-rich microbiota and *C. scindens* strains. Similarly, BAs derived from *Clostridia*-rich microbiota were found to reduce SHP expression in hepatocytes. But interestingly, the *in vitro* results could not be replicated *in vivo* by BA intervention in mice. Whether and how BAs affect hepatic FXR/SHP signaling in humans need to be further systematically investigated. Collectively, as shown in the schematic diagram (Figure. 5E), we propose that the *Clostridia*-rich microbiota result in a specific composition of BAs that attenuate intestinal FXR/FGF19 signaling, thus elevating hepatic synthesis and fecal excretion.

It is well-known that there is an intestinal crosstalk between gut bacteria and BAs.^19^ We clarified inhibitory effects of *Clostridia*-rich microbiota on intestinal BA feedback control, but the effects of BAs on gut microbiota were not involved. In fact, different BA components, in turn, show diverse effects on specific bacterial survival or transforming actions. The presence of TCA, GCA and CA cannot influence the growth of *Clostridium difficile* (*C. difficile*) *in vitro*.^57^ Contrarily, secondary BAs, like DCA, LCA and UDCA, inhibit the germination and/or growth of *C. difficile*,^58–60^ of which DCA and LCA are also proved to enhance inhibitory effects of tryptophan-derived antibiotics secreted by 7α-dehydroxylating species (*C. scindens* and *C. sordellii*) against *C. difficile*.^61^ Different from *in vitro* studies, we did identify the existence of *C. difficile* in fecal communities of all recruits but without difference among groups, suggesting the 7-dehydroxylating species-driven antibacterial effect may not be significantly presented unless *C. difficile* dominates gut ecosystem. Recently, Devendran, *et al.*^62^ revealed that CA significantly elevated expressions of *bai* genes and activity of 7-dehydroxylation for *C. scindens*, but DCA showed no effect on its 7-dehydroxylation, indicating that BA biotransformation of *C. scindens* can be stimulated in presences of abundant primary free BAs. Such findings suggest that increased abundances of *C. scindens* and *bai* genes identified in BA^+^IBS-D patients may also result from the higher contents of CDCA and CA in the lumens. Based on existing evidences from other groups and ours, the *Clostridia*-rich/BA-excessive signature is believed as the consequence of a crosstalk between gut microbiota and BA metabolism in IBS-D.

One limitation of this study is that the linkage between *Clostridia*-rich microbiota and enhanced BA synthesis/excretion need to be further verified in longitudinal studies, although standardization of sampling has been conducted between individuals. Another limitation is that patients with type I and III BAD were not included, whether the microbiota-driven mechanism described here is only appeared in primary BAD (type II) needs to be further investigated.

In conclusion, by using integrative metagenomic, metabolomic and mechanistic approaches, we found that a *Clostridia*-rich microbiota can enhance BA excretion. We conclude that this occurs through a reduction in FGF19/15 release, resulting in loss of feedback inhibition of hepatic BA synthesis. Our study suggests that the double signature concept, *Clostridia*-rich microbiota with excessive BA synthesis, could be the basis for more precise diagnosis and symptom management for IBS-D. More broadly, this approach i.e., assessing the gut microbial signature together with its metabolic features, could be applied in microbiota-associated physiological, pathophysiological and therapeutic research, not just for IBS, but for all gut microbiota-related diseases.

## Acknowledgments

We thank Prof. Zongwei Cai from department of Chemistry, Hong Kong Baptist University, for supplying bile acid standards. We also thank all the patients and healthy volunteers who donated specimens for this study.

## Author contributions

WJ, ZXB and XDF jointly designed this study. WJ and ZXB supervised the study and revised manuscript. HEN and APL contributed project discussion and gave comments for this study. LZ, WY, YC and FJH drafted manuscript. LZ, WY, LL, LXZ, ZWN and CYL collected clinical samples and performed tests of clinical samples. LZ, WY and LL performed experiments of bacterial manipulation. WY and FJH tested the effects of BAs on FXR signaling in vivo and in vitro. ZXR and DFC provided supporting for animal experiments and protein assay. TH provided guidance in data analysis and R script. LLZ, WCL, CWC, ZY, XZ and ZXB recruited IBS patients. YC, LJH, QWQ, XXS, RYH and SLS are responsible for fecal DNA extraction, sequencing and metagenomic data analysis.

## Data availability

Fecal metagenomic sequencing reads can be downloaded from CNGB Nucleotide Sequence Archive (accession number CNP0000334), and other data that support the findings of this study are available from the corresponding author upon reasonable request.

## Ethics approval

The clinical study received approval from the Committee on the Use of Human & Animal Subjects in Teaching & Research in Hong Kong Baptist University (HASC/15-16/0300 and HASC/16-17/0027). Animal experiments followed the Animals Ordinance guidelines, Department of Health in Hong Kong and reporting of *in vivo* experiments also followed ARRIVE guidelines.

## SUPPLEMENTARY MATERIALS

Supplementary methods

Supplementary figures include

Figure. S1. Alteration of serum bile acid profile in IBS-D patients.

Figure. S2. Association of *Clostridial* abundances with BA synthetic level in IBS-D patients. Figure. S3. Changed luminal BA transformation associated with BA synthetic regulation in mouse with transplantation of IBS-D fecal microbiota.

Figure. S4. The effects of *Clostridium* species on ileal BA feedback control *in vivo* and hepatic synthetase *in vitro*.

Figure. S5. Regulatory effects of *Clostridia*-derived BA individuals on hepatic FXR signaling.

Supplementary tables include

Table S1. The MRM transitions and MS parameters for BA analytes and IS. Table S2. The demographics of each donors for FMT experiments.

Table S3. Spearman’s correlation between fecal total BA level and other clinical indexes in included subjects.

Table S4. Data production, quality control, gene prediction resulted from fecal metagenomic sequencing analysis.

Table S5-7. Relative abundances of differential phyla, genera and species in IBS-D patients. Table S8. Relative abundances of KOs related to bile acid biotransformation.

Table S9. Oligonucleotide primers for quantitative RT-PCR analysis.

