## Supplemental methods and figures for "A *Clostridia*-rich microbiota contributes to increased excretion of bile acids in diarrhea-predominant irritable bowel syndrome"

**Supplementary materials include**

Supplementary methods

Supplementary figures include

Figure. S1. Alteration of serum bile acid profile in IBS-D patients.

Figure. S2. Association of *Clostridial* abundances with BA synthetic level in IBS-D patients.

Figure. S3. Changed luminal BA transformation associated with BA synthetic regulation in mouse with transplantation of IBS-D fecal microbiota.

Figure. S4. The effects of *Clostridium* species on ileal BA feedback control *in vivo* and hepatic synthetase *in vitro*.

Figure. S5. Regulatory effects of *Clostridia*-derived BA individuals on hepatic FXR signaling.

### SUPPLEMENTARY METHODS

***Subject recruitment and sampling***

IBS-D patients were recruited through the advertisements or press released in local newspaper. The detail criteria used for patient recruitment are shown as follows:

*Inclusion criteria:* IBS-D patients were recruited on if they fulfilled the following criteria: 1) meet of Rome IV criteria, including recurrent abdominal pain on average at least 1d/week in the last 3 months; 2) appearance of stool form with at least 25% of loose or watery stools and fewer than 25% of hard stools based on Stool Bristol Score; 3) IBS Symptom Severity Scale (IBS-SSS) over than 75 points at baseline; 4) age of 18 to 65 years; 5) normal colonic evaluation with 5 years by examination of colonoscopy or barium enema; 6) Written informed consent.

*Exclusion criteria:* Patients were excluded if they have one or more of follows: 1) pregnancy or breast-feeding; 2) medical history of inflammatory bowel diseases, carbohydrate malabsorption, hormonal disorder, known allergies to food additives, and/or any other serious diseases; 3) surgical histories of gallbladder removal, GI tract, and cerebral cranium; 4) having parasitic infections; 5) having suicidal ideas or attempts or aggressive behavior; 6) use of medications known to influence gastrointestinal function, blood pressure and fat.

The sample size of matched healthy controls was expected referred to the around 30% of pooled prevalence of excess total BA excretion in IBS-D population. The recruiting criteria for healthy population is shown as follows:

*Inclusion criteria*: 1) age of 18–65 years (inclusive); 2) no medical history of metabolic disorders, cardiovascular diseases, neurodegenerative diseases, and gastrointestinal diseases; 4) normal hepatic, renal, and bowel functions within 3 years; 4) no drug taken history for chronic diseases, metabolic diseases, cardiovascular and cerebrovascular diseases, psychiatric illness, disease of immune system, and other serious diseases within 1 year; 5) written informed consent.

*Exclusion criteria*: 1) pregnancy or breast-feeding; 2) surgical histories of gallbladder removal, GI tract, and cerebral cranium; 3) having parasitic infections; 4) use of medications known to influence gastrointestinal transit, blood pressure and fat.

A questionnaire of IBS symptoms contained IBS-SSS, defecation frequency within a day and Bristol Stool Form Scale was required to complete when the interview with physicians. Fecal consistency of individuals was assessed by the score of the Bristol Stool Form Scale. Meanwhile, another dietary questionnaire that recorded complete diet information and dietary habits within past three months was also required to be completed upon subject recruitment and, those with specific dietary habits, such as alcohol consumption or a completely vegetable-based diet, were excluded as well. Participants were instructed to provide fasting blood samples and morning first feces for analyzing 1) serum biochemical indices and stool culture test (conducted by Chan & Hou Medical Laboratories Ltd, Hong Kong); 2) bile acid (BA) profiles and metagenomics. They were also required to stop using antibiotics, probiotics, prebiotics and other microbiota-related supplements at least one month before sampling.

***Targeted bile acid profiling based on UPLC/MS***

*Chemicals and reagents*

A total of 36 BA metabolites, including cholic acid (CA), ω-muricholic acid (ωMCA), α-muricholic acid (αMCA), β-muricholic acid (βMCA), hyocholic acid (HCA), hyodeoxycholic acid (HDCA), chenodeoxycholic acid (CDCA), ursodeoxycholic acid (UDCA), deoxycholic acid (DCA), isodeoxycholic acid (IsoDCA), isolithocholic acid (IsoLCA), 7-ketodeoxycholic acid (7-KDCA), 6-ketolithocholic acid (6-KLCA), 7-ketolithocholic acid (7-KLCA), 12-ketolithocholic acid (12-KLCA), dehydrocholic acid (DHCA), lithocholic acid (LCA), allo-cholic acid (ACA), 23-nordeoxycholic acid (23-NDCA), glycocholic acid (GCA), glycochenodeoxycholic acid (GCDCA), glycodeoxycholic acid (GDCA), glycolithocholic acid (GLCA), glycodehydrocholic acid (GDHCA), glycohyodeoxycholic acid (GHDCA), glycohyocholic acid (GHCA), glycoursodeoxycholic acid (GUDCA), taurocholic acid (TCA), taurodeoxycholic acid (TDCA), tauroursodeoxycholic acid (TUDCA), taurohyocholic acid (THCA), taurohyodexoycholic acid (THDCA), taurochenodeoxycholic acid (TCDCA), taurolithocholic acid (TLCA) were purchased from Sigma-Aldrich (St. Louis, MO, USA). Tauro-α-muricholic acid (TαMCA) and Tauro-β-muricholic acid (TβMCA) were purchased from Santa Cruz Biotechnology (Santa Cruz, CA, USA). An isotopic BA deoxycholic acid-2,2,4,4-d4 (DCA-d4), served as internal standard and was obtained from CDN isotopes (Pointe-Claire, Quebec, Canada). HPLC grade organic reagents for mass spectrometric analysis were purchased from Sigma-Aldrich (St. Louis, MO, USA).

*Stock solution and calibration curve preparation*

All BA chemical standards were separately dissolved in methanol as stock solution with a concentration of 5 mg/ml. A mixed stock solution was obtained after mixing individual standard stock solution. Diluting stock solutions in methanol, the working solution were prepared at a series concentration of 0.020, 0.102, 0.512, 2.56, 12.8, 64, 320, 1600, 8000, and 40000 ng/ml for individual BAs, while at a series concentration of 0.064, 0.32, 1.6, 8, 40, 200, 1000, and 5000 ng/ml for serum C4. The standard curves and regression coefficients were gained based on IS adjustment. The signals of each BA metabolites were found in individual measured ranges.

*UPLC/TQ-MS condition*

An ultra-high-performance liquid chromatography (Agilent UHPLC 1290, USA) coupled with a triple-quadrupole mass spectrometer (Agilent QQQ-MS 6438, USA) was applied for bile acid analysis. Though a single 26-min acquisition with positive/negative ion switching, bile acid metabolites (under ESI-) and C4 (under ESI+) were simultaneously quantified in multiple reaction monitoring (MRM) mode. Sample injection and flow rate were set at 2 μL and 0.35 ml/min for each sample, respectively. Bile acid metabolites were separated using a ACQUITY BEH C18 column (1.7 μm, 100mm × 2.1 mm) with a linear gradient of 0.1% formic acid (FA) in water (A) and 0.1% FA in acetonitrile (B). The gradient program was: 25% to 40% B for the first 6 min, 40% to 70% B for 14 min, 70% to 100% B for 0.1 min, held at 100% B for 2.9 min, then re-equilibration at 25% B for 0.1 min, and held at 25% B for 2.9 min. The column temperature was maintained at 45℃. The capillary voltage of mass spectrometer was 3.5 kV and 4 kV in positive and negative modes. The acquisition data was analyzed using Agilent MassHunter Workstation Software for peak integration, calibration equations and quantification of individual BAs.

***Measurement of bacterial BA-transforming activities***

The BA-transforming activities of gut bacteria isolated from human feces or mouse cecal contents were assessed *in vitro* by quantifying the BA products and substrates for each transforming action using LC/MS. The concentration of supplemented BA substrates was referred to the level of total BAs in feces of human beings. In details, mixed bacteria were individually extracted from human feces using 10-fold sterile PBS with differential centrifugation. A volume of 1mL of bacterial reaction mixture, containing bacterial extracts (diluted as 0.1 of the final OD600 value), BHI medium and 5 mM BA substrate (GCDCA, CDCA or CA), were prepared and shaken continuously at 37°C for 24 hours. Then, BA metabolites were extracted from the culture medium using 3-fold methanol for LC/MS-based quantitative analysis. The ratio of CDCA to GCDCA represented the level of bacterial deconjugation, the ratio of UDCA to CDCA and DCA to CA were used for evaluating bacterial 7-HSDH and 7α-dehydroxylating levels, respectively. For mouse BA-transforming activity analysis, Mixed bacteria were isolated from cecal contents, and the bacterial reaction mixture with 5 mM BA substrate (TCA, CDCA or CA) were prepared as mentioned above. The level of bacterial deconjugation was evaluated by the ratio of TCA to CA, while bacterial 7-

HSDH and 7α-dehydroxylating levels were tested with the ratio of UDCA to CDCA and the ratio of DCA to CA, respectively.

***Quantitative real-time PCR analysis of bacterial and host genes***

Total DNA was extracted from cecal contents (100 mg) of FMT or *Clostridium*-treated mice for specific bacteria analysis using real-time PCR detection. Briefly, a reaction mixture consisting of DNA template (50ng), 0.5μM each DNA oligonucleotide primers, and SYBR Green PCR Master Mix (2X) was prepared and amplified using a real-time PCR cycler (ViiA$^{TM}$ 7Dx Instrument, Applied Biosystems, Foster city, CA, USA). The relative levels of each bacterium from each DNA samples was normalized to the total bacterial expression.

Moreover, total RNA samples were individually isolated from the hepatic and ileum tissues of experimental mice using TissueLyzer (Qiagen, Hilden, Germany) with Trizol reagent (Invitrogen, Carlsbad, CA, USA). The cDNAs were produced from RNA samples using the SuperScript® First-Strand synthesis system (Invitrogen, Carlsbad, CA, USA). Quantitative real-time PCR detection was performed on the ViiA™ 7 Real-Time PCR System, and analysis of target gene expression was processed with the $\Delta\Delta C_T$ method. Oligonucleotide primers for bacteria and host genes related to BA metabolism are summarized in **Table S9**.

***Immunoblot analysis of proteins in mouse tissues, hepatic and enteric cells***

Proteins were extracted from mouse tissue (50mg) using RIPA buffer, loaded on a 10% SDS-PAGE gel and blotted onto a PVDF membrane using the Trans-Blot® TurboTM Transfer system (Bio-Rad Laboratories). The membranes were blocked with Odyssey Blocking buffer (P/N 927-50000), and then incubated with primary rabbit anti-actin (1: 5000, ab8227), mouse anti-CYP7A1 (1:500, sc-293193) or mouse anti-FGF15 (1:500, sc-514647) overnight at 4 °C. After incubation with anti-rabbit and anti-mouse antibodies conjugated with IRDye 680RD or 800CW, membranes were scanned on the Odyssey Infrared Imaging System (LICOR Biosciences) with Image Studio Lite Software (v5.2, LI-COR Biosciences).

Moreover, proteins related to BA synthetic regulation were separately analyzed in total protein extracts from human cells (FXR, SHP, CYP7A1, CYP8B1 for hepatocytes; FXR and FGF19 for enterocytes). Accordingly, primary rabbit anti-actin (1: 5000, ab8227), goat anti-FXR (1: 1000, ab51970), mouse anti-SHP (1:500, sc-271511), mouse anti-CYP7A1 (1:500, sc-293193), mouse anti-FGF19 (1:500, sc-390621) were used for the performance of Western Blots.

### SUPPLEMENTARY FIGURES

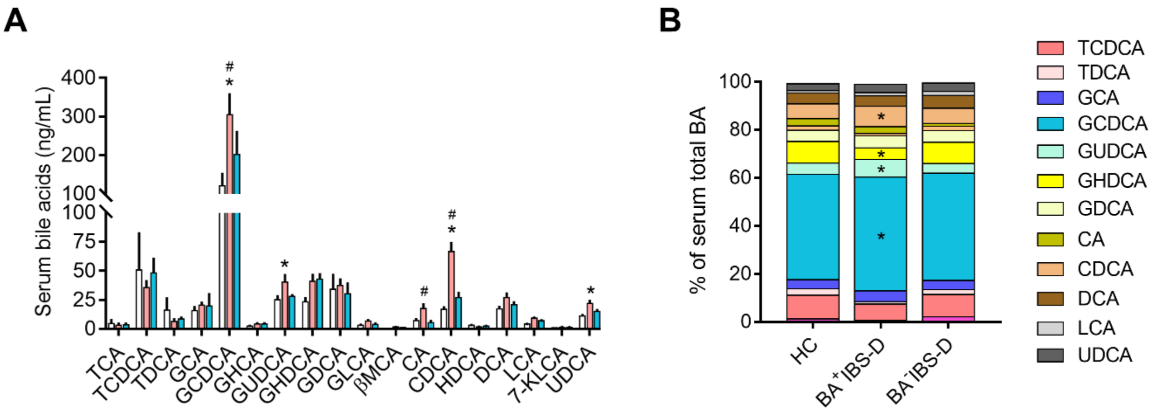


**Figure. S1.** Alteration of serum bile acid profile in IBS-D patients. (A) Absolute contents of serum dominant BA metabolites. (B) Proportions of serum dominant BAs. Only BAs with over 1% of total BA are shown in the legend. Serum BA individuals were analyzed by LC/MS. Comparison of BA metabolites was analyzed by the Kruskal-Wallis test. Statistical significance is expressed by *, $p$<0.05 compared with the HC group; #, $p$<0.05 compared with the BA$^-$IBS-D group.

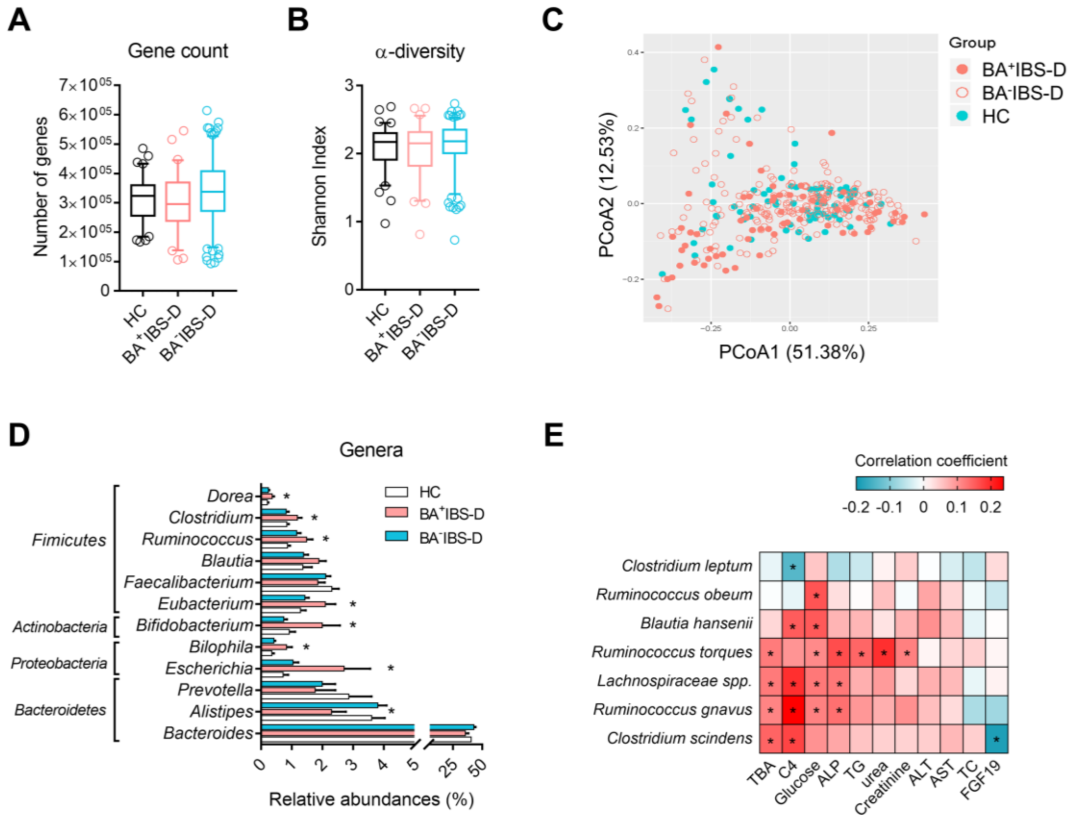


**Figure. S2.** Association of *Clostridial* abundances with BA synthetic level in IBS-D patients. (A) The total gene count obtained from the metagenomic dataset of human fecal samples. (B) Microbial α-diversity measured by the Shannon index. (C) Principle component analysis of human microbial communities based on the Genus-level Bray-Curtis dissimilarity. (D) The relative abundances of dominant genera identified from human fecal microbiomes. (E) Relationships between abundances of BA-transforming *Clostridial* species and clinical biochemical indexes resulted from Spearman's correlation analysis. Differential taxa among three groups were analyzed with the Benjamin-Hochberg method, statistical significance is expressed by *, $p$<0.05; **, $p$<0.001 compared with the HC group. Spearman's correlation was performed in GraphPad Prism 7, and statistical significance is set as *, $p < 0.05$.

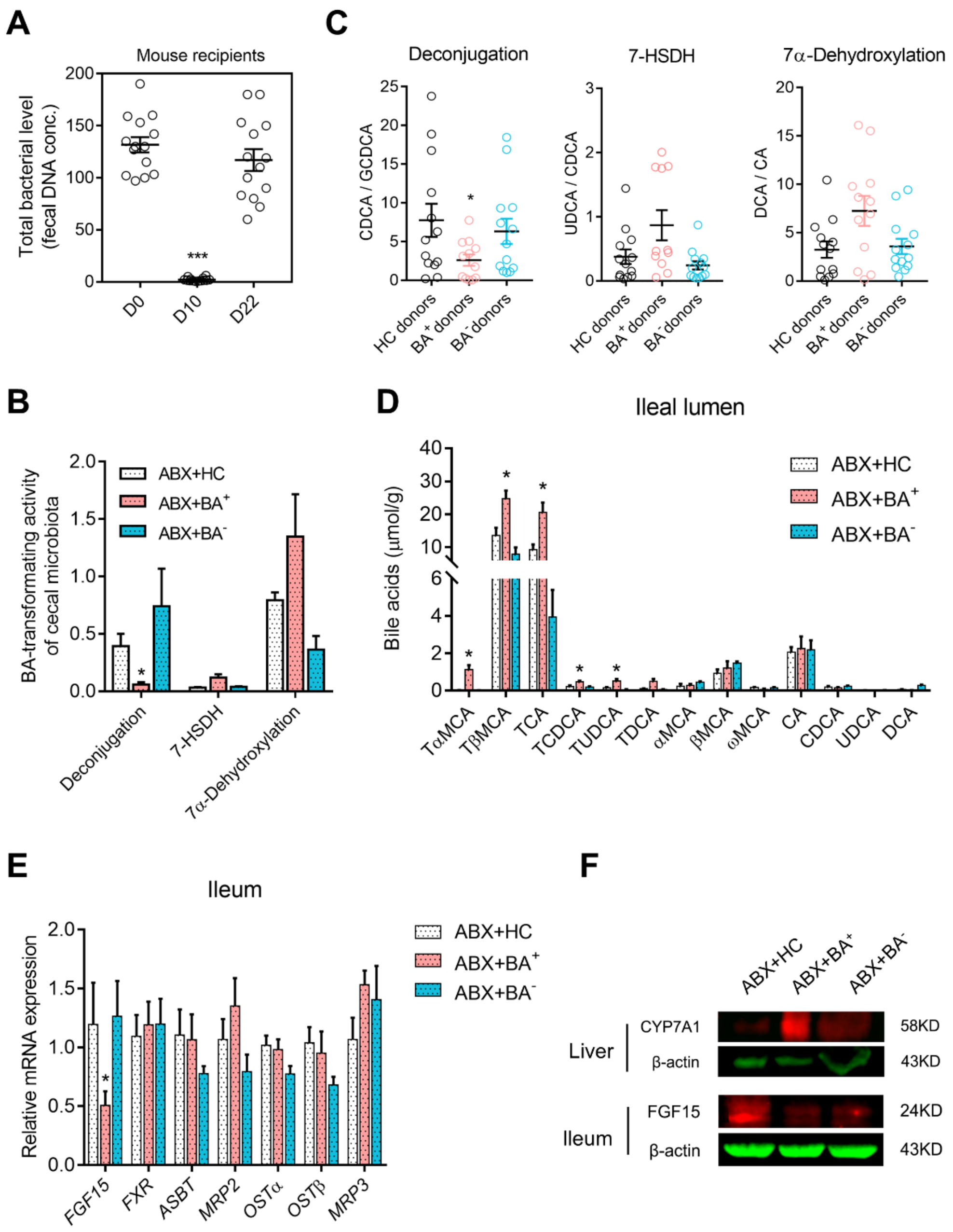


**Figure. S3.** Changed luminal BA transformation associated with BA synthetic regulation in mouse with transplantation of IBS-D fecal microbiota. (A) Dynamic alteration of cecal total bacterial counts throughout the FMT experiment. (B, C) Bacterial BA-transforming levels of donors and mouse recipients. Each BA-transforming activity was measured by the ratio of the BA product to substrate. (D) The BA profile of the ileal contents in mouse recipients. (E) Relative gene expression of ileal proteins related to BA feedback and transport in mouse recipients. (F) Protein expressions of hepatic

CYP7A1 and ileal FGF19. Differential BA-related bacteria, metabolites and genes among groups of either donors or recipients were evaluated by the Kruskal-Wallis test. Statistical significance is expressed by *, $p<0.05$; ***, $p<0.005$ compared with individual control group.

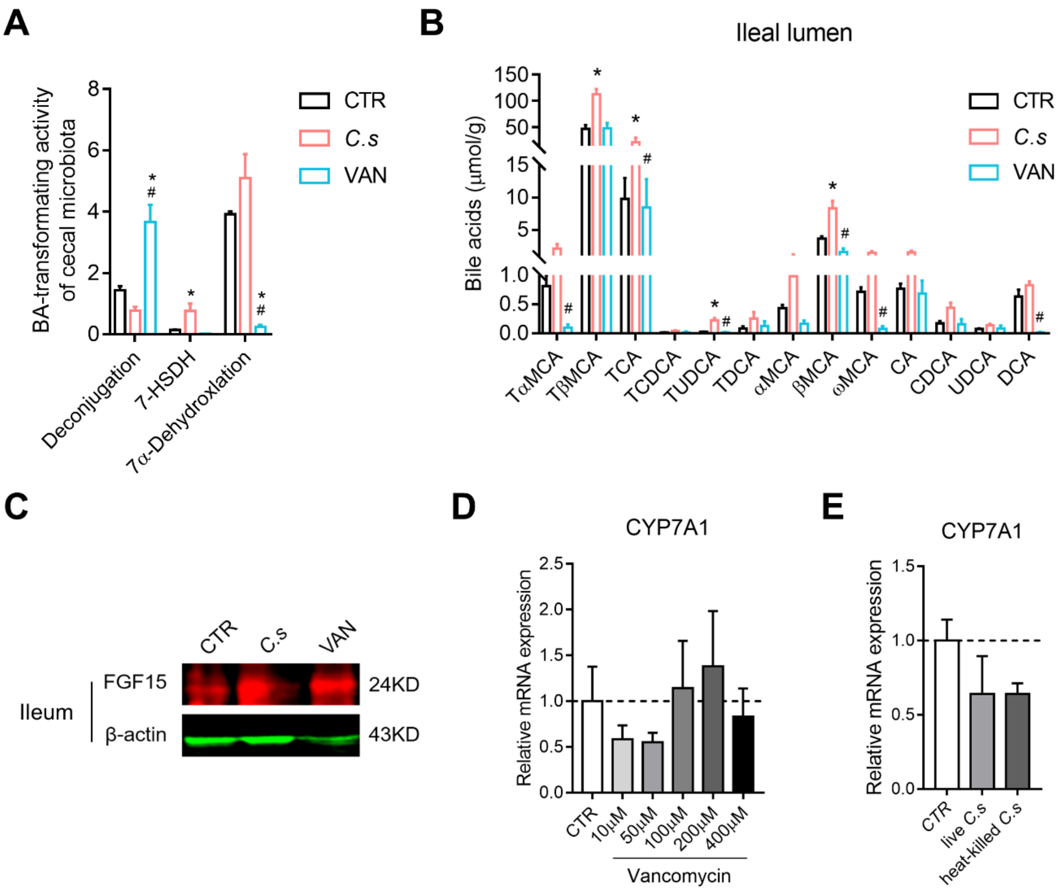


**Figure. S4.** The effects of *Clostridium* species on ileal BA feedback control *in vivo* and hepatic synthetase *in vitro*. (A) Cecal microbial BA-transforming levels in mice with manipulation of *Clostridium* species. (B) Luminal BA profile of mouse ileum. (C) Protein expression of ileal FGF15. (D, E) Gene expressions of CYP7A1 when culturing hepatocytes in presence of vancomycin, live or heat-killed *C. scindens*. Differential BA-related results among three groups were evaluated by the Kruskal-Wallis test. Statistical significance is expressed by *, $p<0.05$ compared with the control group; #, $p<0.05$ compared with the *C. scindens* group.

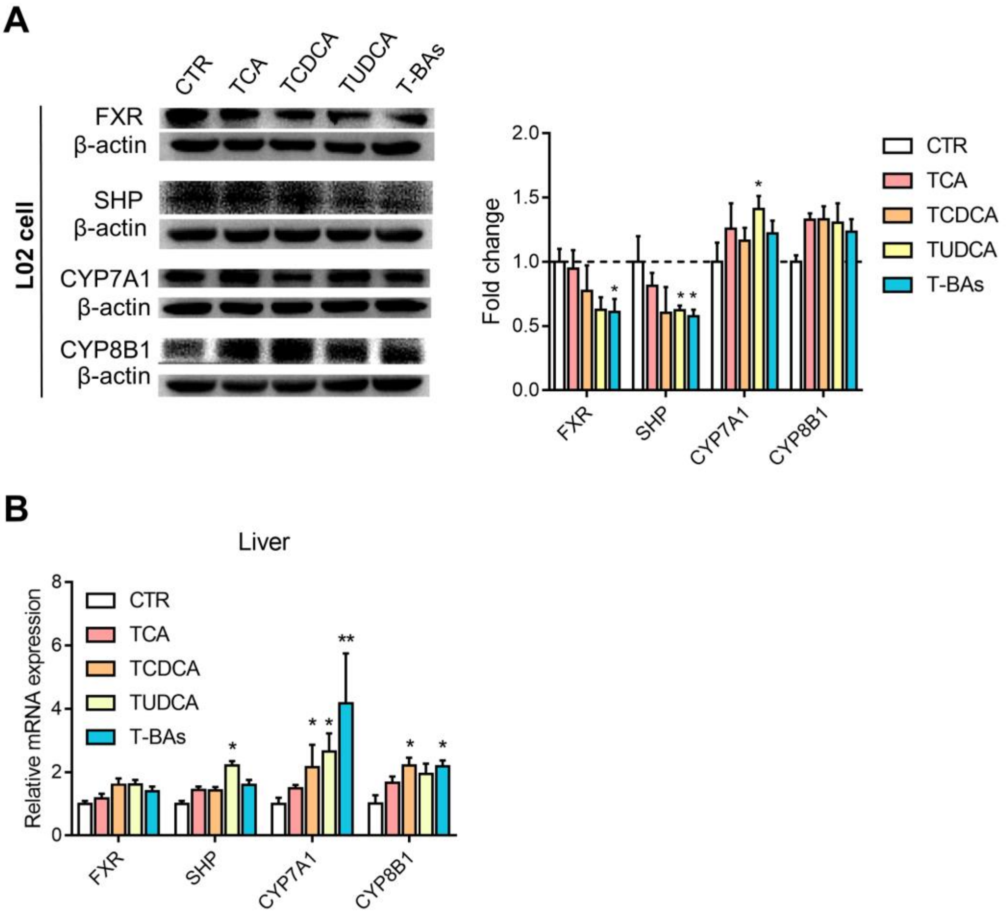


**Figure. S5.** Regulatory effects of *Clostridia*-derived BA individuals on hepatic FXR signaling. (A) Effects of taurine-conjugated BAs on protein expressions of FXR, SHP, CYP7A1 and CYP8B1 in hepatocytes. (B) Gene expressions of FXR, SHP, CYP7A1 and CYP8B1 in the liver of mice subjected to treatment of taurine-conjugated BAs. Differential genes or proteins were analyzed with the Kruskal-Wallis test, and statistical significance is expressed by *, $p<0.05$; **, $p<0.01$ compared with the control group.
